# The CARD8 inflammasome drives CD4^+^ T-cell depletion in HIV-1 infection

**DOI:** 10.1101/2023.03.18.533283

**Authors:** Qiankun Wang, Liang Shan

## Abstract

CD4^+^ T-cell depletion is the root cause of acquired immunodeficiency syndrome (AIDS). How HIV depletes CD4^+^ T cells in humans remains unknown because vast majority of the dying CD4^+^ T cells in patients are uninfected. Burgeoning evidence supports the hypothesis that non-productive HIV-1 infection triggers CD4^+^ T-cell depletion in the course of pathogenic HIV and SIV infections. Here, we report that the CARD8 inflammasome is activated immediately after HIV-1 entry by the viral protease encapsulated in the incoming HIV-1 particles. Sensing of HIV-1 protease activity by the CARD8 inflammasome leads to rapid pyroptosis of quiescent CD4^+^ T cells without productive viral infection. In humanized mice reconstituted with a CARD8-deficient immune system, CD4^+^ T-cell loss is delayed despite increased levels of HIV-1 replication. Our study suggests that the CARD8 inflammasome drives CD4^+^ T-cell depletion and disease progression through rapid sensing of HIV-1 particles.

## Introduction

The hallmark of human immunodeficiency virus type 1 (HIV-1) pathogenesis is a progressive depletion of CD4^+^ T-cell populations, the underlying cause of increased susceptibility to opportunistic infections and progression to acquired immunodeficiency syndrome (AIDS). The mechanism through which HIV depletes CD4^+^ T cells in humans has been the subject of intense research for decades. Although HIV-1 proteins mediate cytopathic effects, CD4^+^ T cell depletion is not confined to virus-infected cells, because only around 1 in 10^4^ peripheral blood mononuclear cells (PBMCs) or up to 1% CD4^+^ T cells were infected in untreated individuals ^1-5^. Notably, only cells positive for HIV-1 RNA, DNA, or protein were deemed infected cells in these studies, hereinafter referred to as productive infection. In productively infected cells, HIV-1 completes all steps in its life cycle and produces viral RNA and proteins. However, viral RNA was rarely observed in the dying CD4^+^ T cells during HIV-1 infection ^6^. Since other studies show that CD4 depletion required HIV-1 co-receptor expression ^7,8^, it is likely that the co-receptor-mediated viral particle and target cell interaction or viral entry is a prerequisite for cell death, but productive HIV-1 infection is not required. Similarly, simian immunodeficiency virus (SIV) infections drive rapid depletion of non-productively infected CCR5-expressing CD4^+^ T cells in Asian non-human primates (NHPs) ^6,9,10^. By contrast, SIV infections of African NHPs do not lead to systemic CD4^+^ T-cell loss, even though the virus is equally cytopathic in productively infected CD4^+^ T cells and plasma viral loads are comparable between different NHPs ^11^.

In Asian NHPs, rapid inflammasome activation was observed at the site of SIV inoculation and the sites of distal virus spread ^12^. More importantly, caspase-1(CASP1)-dependent pyroptosis was the dominant mechanism responsible for the rapid CD4 depletion by SIV, whereas other programmed cell death mechanisms contribute minimally ^13^. In humans, CASP1 activation and pyroptotic cell death were observed in uninfected CD4^+^ T cells from viremic individuals ^14^. Taken together, these results suggest that HIV-1 cytopathic effects in productively infected cells are not the major cause of CD4^+^ T-cell destruction in humans and viral infection drives co-receptor-dependent loss of uninfected cells. However, how HIV-1 engages its co-receptors to trigger inflammasome activation in CD4^+^ T cells remains unknown. Since most inflammasomes have been identified and characterized in myeloid cells, their roles in human CD4^+^ T cells are not well defined. We recently reported that Caspase recruitment domain-containing protein 8 (CARD8) can detect HIV-1 protease activity and mediates assembly of the inflammasome complex ^15^, but its physiological role in HIV-1 infection and pathogenesis is unclear. The CARD8 C-terminus contains a “function-to-find” domain (FIIND), followed by a CARD domain. CARD8 undergoes autoproteolytic processing in the FIIND domain, generating the N-terminal ZU5 and C-terminal UPA-CARD fragments that remain associated noncovalently between the F296 and S297 positions ^16^. CARD8 can be activated by direct proteolysis of its N terminus by viral proteases ^15,17^, which results in an unstable neo–N terminus targeted for proteasome degradation. Because of the noncovalent bond, the bioactive UPA-CARD subunit is liberated and initiates CASP1-dependent inflammasome assembly. The ability of the CARD8 inflammasome to drive pyroptotic cell death has been demonstrated in different lineages of immune cells ^15,18-20^. In this study, we aimed to determine whether CARD8 is responsible for HIV-1-induced CD4 depletion, and if so, how CARD8 is activated during the natural course of viral infection to drive rapid CD4^+^ T-cell loss.

## Results

### HIV-1 entry induces rapid CD4^+^ T-cell loss through the CARD8 inflammasome

To determine how HIV-1 induces rapid loss of CD4^+^ T cells, activated CD4^+^ T cells pre-infected with HIV-1 were co-cultured with autologous peripheral blood or tonsil mononuclear cells (PBMCs or ToMCs) in the presence of antiretroviral drugs (ARVs). The percentage of CD4^+^ and CD8^+^ T cells was measured by flow cytometry within six hours after co-culture, allowing us to examine cell death immediately after viral entry **(Fig. 1a)**. The CD4 to CD8 ratio was reduced from 2.4 to 0.6 when co-cultured with CD4^+^ T cells pre-infected with HIV_NL4-3_. Inhibitors AMD3100 and T20 that block viral entry completely abolished CD4^+^ T-cell loss despite the presence of viral spreading cells, whereas blocking viral reverse transcription by tenofovir (TFV) or integration by raltegravir (RAL) did not prevent CD4^+^ T-cell depletion **(Fig. 1b-d)**. These results suggested that cell death occurred post-viral entry but before reverse transcription. When CD4^+^ T cells pre-infected with the CCR5-tropic HIV_BaL_ were used for the co-culture experiments, a rapid depletion of CCR5^+^ CD4^+^ T cells was observed **(Fig. 1e, f)**, demonstrating that this effect was entry-dependent and was not limited to X4-tropic viruses.

**Fig 1:**
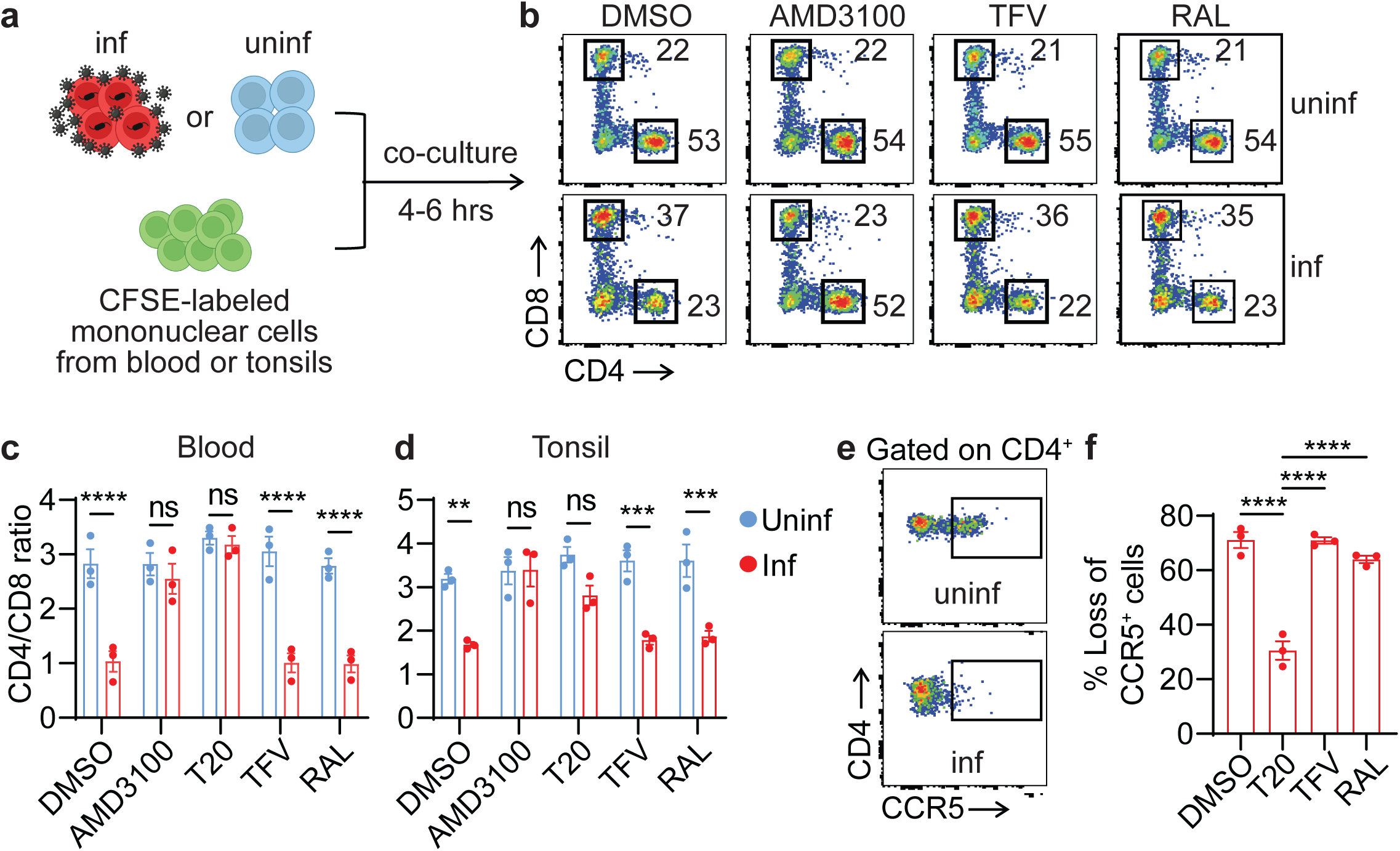
HIV-1 entry triggers rapid loss of CD4^+^ T cells. **a**, The co-culture scheme. Activated blood or tonsillar CD4^+^ T cells were infected with either HIV_NL4-3_ or HIV_BaL_ for three days. Virus-producing cells were then co-cultured at a 1:1 ratio with CFSE-labeled donor-matched unstimulated mononuclear cells from blood or tonsils for four to six hours with the presence of indicated ARVs. **b**, Representative plots were shown from blood cells infected with HIV_NL4-3_. **c, d**, Rapid loss of blood and tonsillar CD4^+^ T cells. Virus-producing cells were infected with HIV_NL4-3_. Three blood samples and three tonsillar samples were used. **e, f**, Rapid loss of CCR5^+^ CD4^+^ T cells. Virus-producing cells were infected with HIV_BaL_. Three blood samples were used. In **c** and **d**, *p* values were calculated using the two-way ANOVA with Šidák’s multiple comparison tests. In **f**, *p* values were calculated using the one-way ANOVA with Dunnett tests. ** *p* < 0.01. *** *p* < 0.001, **** *p* < 0.0001. Error bars show mean values with standard errors of the mean (SEM).

To determine whether the rapid CD4^+^ T-cell destruction post-viral entry was due to CARD8-mediated pyroptosis, we first confirmed that it was prevented by the CASP1-specific inhibitor VX765 and the proteasome inhibitor MG132 **(Fig. 2a, b)**. Next, we modified the co-culture system to test Cas9-edited CD4^+^ T cells **(Fig. 2c)**. We found that *CARD8-, CASP1-*, and *GSDMD*-KO CD4^+^ T cells were resistant to HIV-1 entry-mediated cell killing, whereas NLRP1 and other ASC-dependent inflammasomes were not involved in this process (**Fig. 2d, e**). Compared to the six-hour co-culture, a longer co-culture (48 hours) that completed the viral reverse transcription did not affect the resistance in *CARD8-, CASP1-*, and *GSDMD*-KO CD4^+^ T cells, suggesting that viral reverse transcription products were unable to trigger cell death.

**Fig 2:**
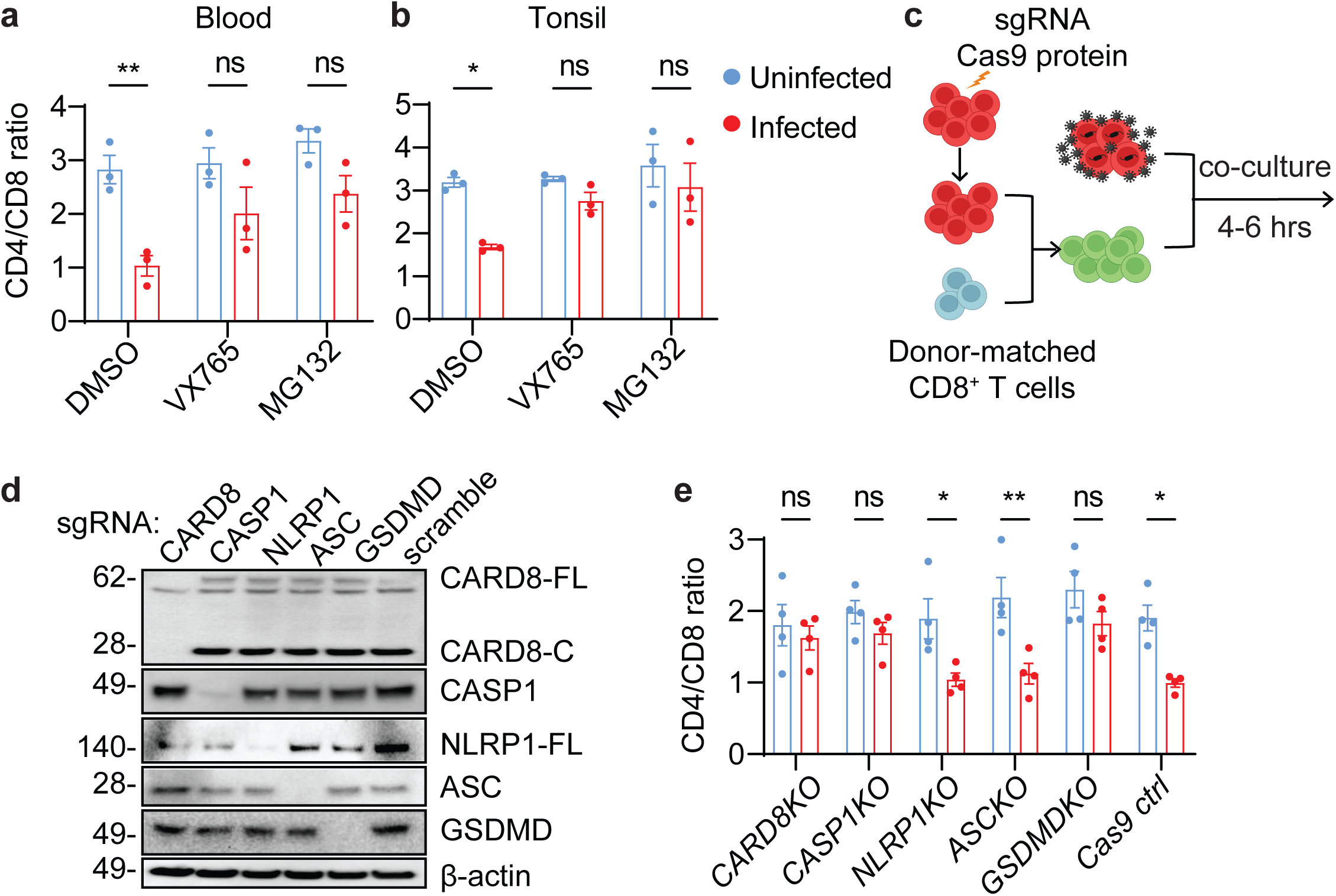
CD4^+^ T-cell depletion by HIV-1 is mediated by the CARD8 inflammasome. **a, b**, Rapid CD4^+^ T-cell death is proteasome- and CASP1-dependent. Unstimulated PBMCs (A) or ToMCs (B) were pretreated with VX765 (50 μM) or MG132 (10 μM) for 30 minutes and then co-cultured with donor-matched uninfected or HIV_NL4-3_-producing CD4^+^ T cells for six hours before flow cytometry analyses. **c**, the co-culture scheme of Cas9-edited unstimulated CD4^+^ T cells and HIV-1-infected autologous CD4^+^ T cells. Unstimulated CD4^+^ T cells electroporated with the indicated gene-specific sgRNA were cultured for three weeks and then mixed with donor-matched CD8^+^ T cells before co-culture with HIV-1-infected CD4^+^ T cells from the same donor. **d, e**, The CARD8 inflammasome is required for the rapid loss of CD4^+^ T cells. Unstimulated CD4^+^ T cells with indicated knockouts were mixed with autologous CD8^+^ T cells at a 2:1 ratio before co-culture with HIV_NL4-3_-producing cells. Cas9 editing efficiency was confirmed in **d**. The immunoblots represent four independent experiments. Four blood samples were used. *p* values were calculated using two-way ANOVA with Šidák’s multiple comparison tests. * *p* < 0.05, ** *p* < 0.01, ns: not significant. Error bars show mean values with SEM from three independent blood donors.

### The CARD8 inflammasome is activated by HIV-1 protease encapsulated in incoming viral particles

Although the co-culture system for viral spreading is physiologically relevant, it requires the use of replication-competent viruses, thus not allowing us to test mutant viruses to determine how the CARD8 inflammasome was activated post HIV-1 entry. Previous studies showed that cell-free viral particles also triggered rapid cell death of CD4^+^ T cells^21^. Therefore, we utilized the cell-free replication-defective HIV-1 reporter virus NL4-3-ΔEnv-EGFP pseudotyped with the NL4-3 envelope for primary CD4^+^ T cells or VSV-G for THP-1 cells to study cell death measured by the LDH release or Zombie plus caspase-1 staining (**Fig. 3a**). Similar to the co-culture experiments, cell death was observed four hours post-exposure to cell-free HIV-1 particles and was blocked by viral entry inhibitors T20 and AMD3100. ARVs that did not block viral entry had no effect on cell death, except for a viral protease inhibitor lopinavir (LPV) (**Fig. 3b-d**). Next, we introduced reverse transcriptase inactive (ΔRT, RT-D110A-D185A) or integrase inactive (ΔIN, IN-D116A) mutations to the NL4-3-ΔEnv-EGFP plasmid, which then generated HIV-1 reporter virus particles carrying either deactivated RT or integrase in their respective viral particles. We found that both mutant viruses induced cell death as efficiently as the control reporter virus (**Fig. 3e**). More importantly, lentiviral particles carrying HIV-1 Gag-Pol triggered comparable levels of cell death as the HIV-1 reporter virus (**Fig. 3f**), suggesting that the viral protease was sufficient to drive CD4^+^ T-cell death. As expected, short-term exposure to HIV-1 particles led to caspase-1 activation and LDH release, which were abrogated by VX765 and MG132 (**Fig. 3g**). Furthermore, deletion of CARD8, CASP1, and GSDMD in CD4^+^ T cells completely abrogated early cell death induced by HIV-1 entry, confirming that HIV-1 infection induces pyroptosis in CD4^+^ T cells through the CARD8 inflammasome (**Fig. 4a**). Dipeptidyl peptidase 9 (DPP9) negatively regulates CARD8 activity by binding to and sequestrating the bioactive CARD8 C-fragment ^22^. Thus, abolishing CARD8 and DPP9 interaction sensitizes the CARD8 inflammasome in cells exposed to HIV-1 particles ^23^. While the DPP9 inhibitor 1G244 alone at ≤1μM did not drive cell death, it greatly enhanced HIV-1 entry triggered pyroptosis of CD4^+^ T cells (**Fig. 4b, c**), further demonstrating that CARD8 is the driver of the post-viral entry cell death.

**Fig 3:**
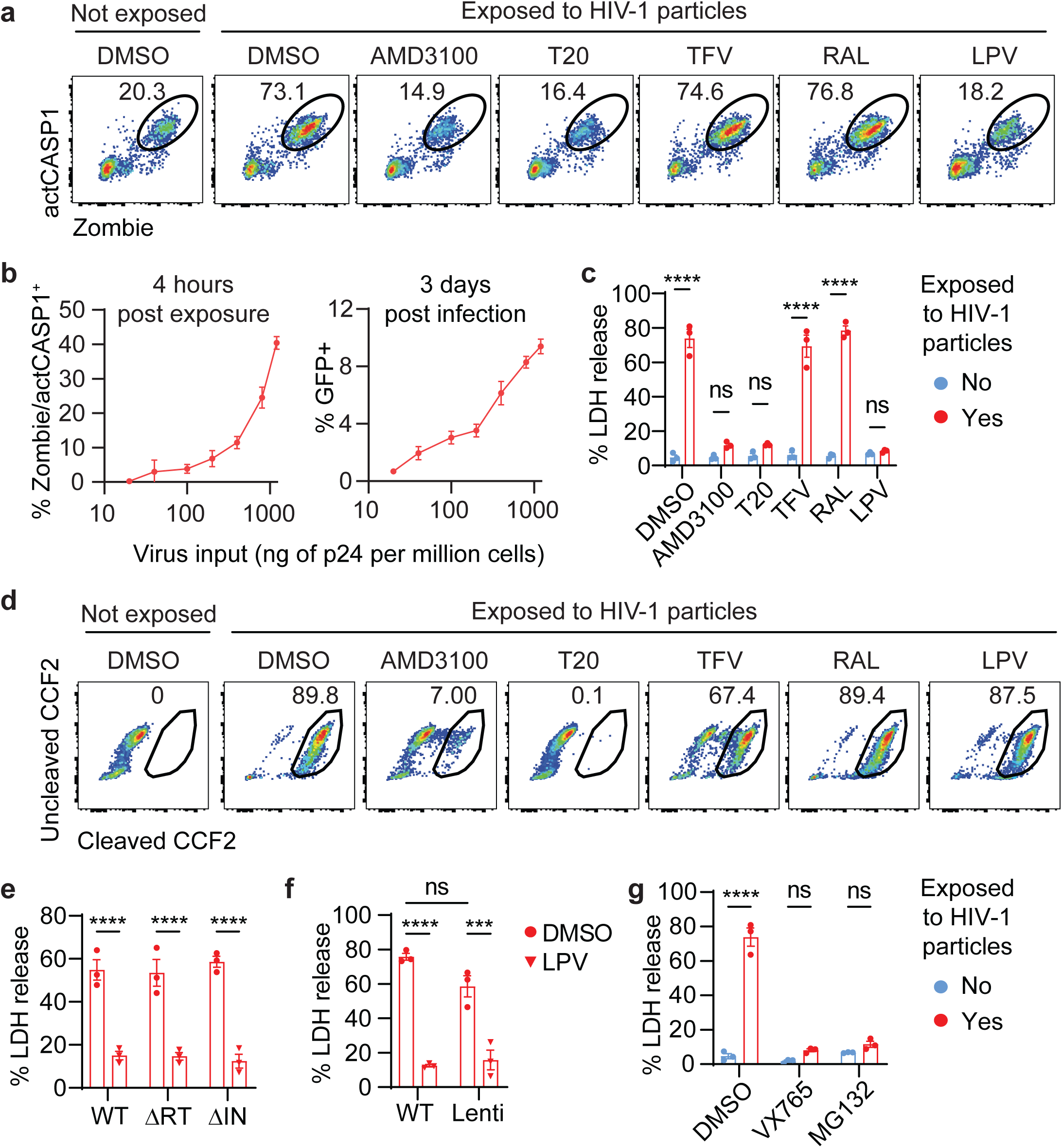
HIV-1 infection induces pyroptosis of CD4^+^ T cells by virion-packaged protease. **a-c**, Rapid loss of CD4^+^ T cells after exposure to HIV-1 particles. Unstimulated blood CD4^+^ T cells were treated with indicated antiretroviral drugs for 30 minutes before being exposed to cell-free HIV-1 reporter virus NL4-3-ΔEnv-EGFP pseudotyped with the NL4-3 envelope. Cell death by LDH release or Zombie/CASP1 staining and viral productive infection by GFP were measured four hours and three days post-infection, respectively. **d**, Quantification of HIV-1 entry. Representative flow cytometry plots of CCF2 substrate cleavage by BlaM in infected blood CD4^+^ T cells. **e, f**, HIV-1 protease encapsulated in incoming viral particles is sufficient to cause CD4^+^ T-cell death. Unstimulated blood CD4^+^ T cells were pre-treated with LPV or DMSO for 30 minutes and then exposed to indicated enzyme inactive reporter HIV-1 (**e**) or lentiviral particles pseudotyped with the NL4-3 envelope (**f**). ΔRT: D110A and D185A. ΔRT: D116A. **g**, Viral protease induced cell death is proteasome- and CASP1-dependent. Unstimulated blood CD4^+^ T cells were pretreated with VX765 (50 μM) or MG132 (10 μM) for 30 minutes before being exposed to cell-free NL4-3-ΔEnv-EGFP pseudotyped with the NL4-3 envelope. *p* values were calculated using the two-way ANOVA with Šidák’s multiple comparison tests. *** *p* < 0.001, **** *p* < 0.0001. The data points are means with SEM and represent three or more independent experiments. N=3.

**Fig 4:**
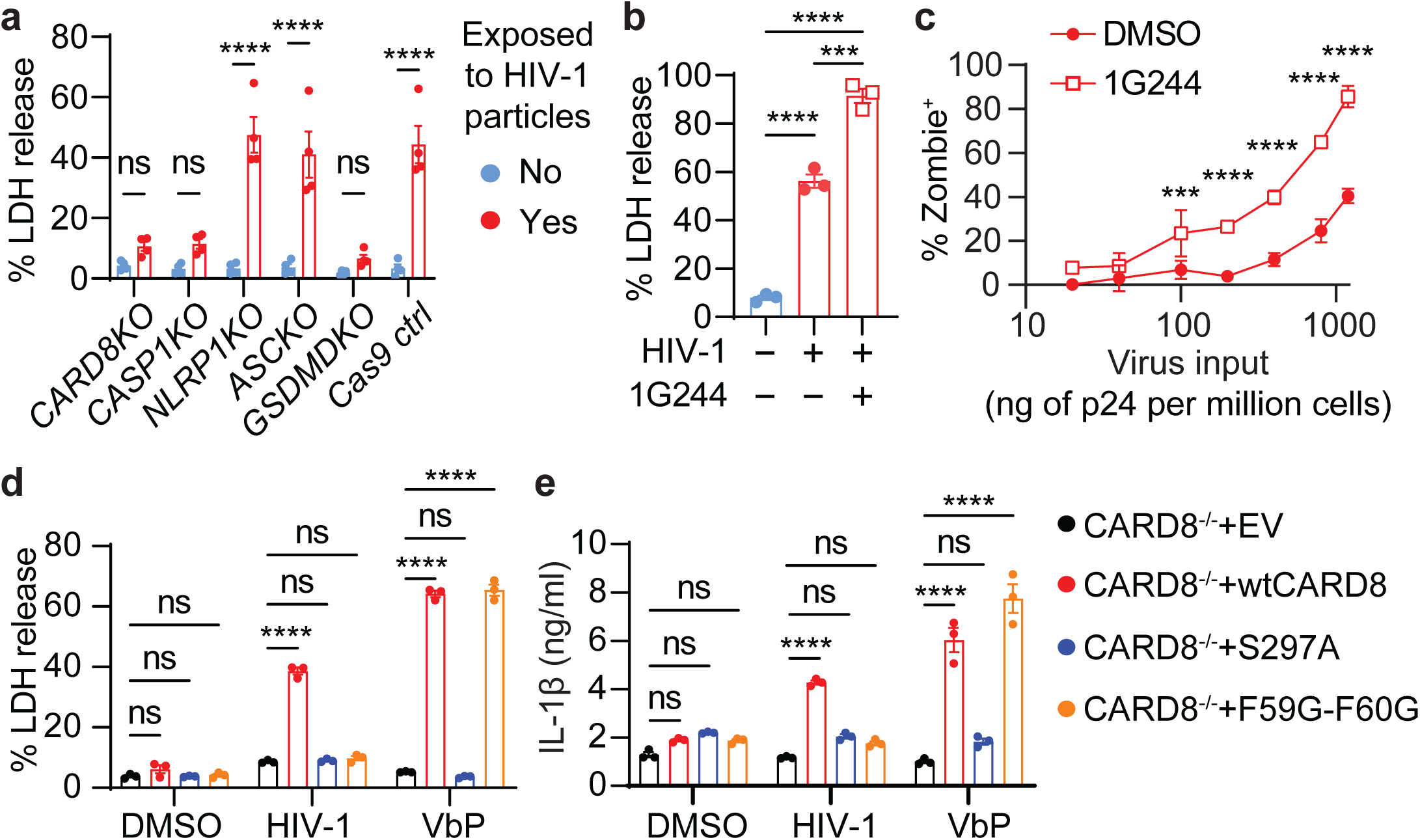
HIV-1 protease encapsulated in incoming viral particles activates the CARD8 inflammasome. **a**, Pyroptosis of CD4^+^ T cells is mediated by the CARD8 inflammasome. Unstimulated CD4^+^ T cells electroporated with the indicated gene-specific sgRNA were cultured for three weeks and then exposed to cell-free HIV-1 reporter virus NL4-3-ΔEnv-EGFP pseudotyped with the NL4-3 envelope for four hours before cell death measurement. **b, c**, 1G244 enhances HIV-1 entry-triggered pyroptosis of CD4^+^ T cells. Unstimulated CD4^+^ T cells were exposed to cell-free virions with or without 1G244 (500 nM) for four hours. **d, e**, Activation of CARD8 inflammasome by HIV-1 in THP-1 cells. *CARD8*-KO THP-1 cells expressing wild-type or mutant CARD8 were exposed to HIV-1 reporter virus NL4-3-ΔEnv-EGFP pseudotyped with the VSVG envelope for six hours before LDH or IL-1β measurement. VbP (5 μM) was used as positive controls. In (**b)**, *p* values were calculated using the one-way ANOVA with Tukey’s multiple comparisons tests. Other *p* values were calculated using the two-way ANOVA with Šidák’s multiple comparison tests. *** *p* < 0.001, **** *p* < 0.0001. The data points are means with SEM and represent three or more independent experiments. N=3.

Our previous study demonstrated that HIV-1 protease cleaves CARD8 between F59 and F60, leading to the formation of an unstable neo-N-terminus for proteasome degradation and subsequent activation of the CARD8 inflammasome ^15^. In this study, we sought to find direct evidence of CARD8 cleavage and activation by HIV-1 particle-derived viral protease. We generated *CARD8*-KO THP-1 cells replete with wild-type (wt), cleavage-deficient (F59G-F60G), or autoprocessing-deficient (S297A) CARD8. Autoprocessing is required for HIV-1 protease-induced CARD8 inflammasome activation, because only the autoprocessed CARD8 can release its bioactive C-terminus from the proteasome complex due to the non-covalent bond. HIV-1 triggered LDH release and IL-1β secretion with the presence of the wtCARD8, whereas both wtCARD8 and the F59G-F60G mutant restored VbP-triggered cell death and IL-1β secretion in *CARD8*^−/−^ THP-1 cells (**Fig. 4d, e**). These results demonstrate that the HIV-1 protease encapsulated in the incoming viral particles is required and sufficient to trigger CARD8-mediated pyroptosis immediately after viral entry.

### The CARD8 inflammasome promotes CD4^+^ T-cell depletion in humanized mice infected with HIV-1

To evaluate the role of the CARD8 inflammasome in CD4^+^ T-cell depletion in vivo, we generated human *CARD8*-KO or *Cas9* control CD34^+^ cells to engraft immunodeficient mice. Deep sequencing analysis showed that 75-90% of human immune cells in mice were CARD8-edited (**Fig. 5a, b**). CARD8-editing did not impede overall human hematopoietic cell reconstitution (**Fig. 5c**) or T cell development (**Fig. 5d**). Next, we infected mice with HIV_BaL_ and measured plasma viral loads and CD4^+^ T cells in different tissues. Mice reconstituted with *CARD8*-KO immune system had higher levels of plasma HIV-1 RNA (**Fig. 5e**), suggesting that CARD8 restricted HIV-1 replication, which was consistent with our in vitro experiments. More importantly, CD4^+^ T-cell loss was delayed in mice with a *CARD8*-KO immune system despite increased levels of viral replication. The frequency and number of CD4^+^ T cells in the tissues were higher in mice with a *CARD8*-KO immune system (**Fig. 5f, g**). Our study in humanized mice demonstrates that the CARD8 inflammasome promotes CD4 depletion during HIV-1 infection.

**Fig 5:**
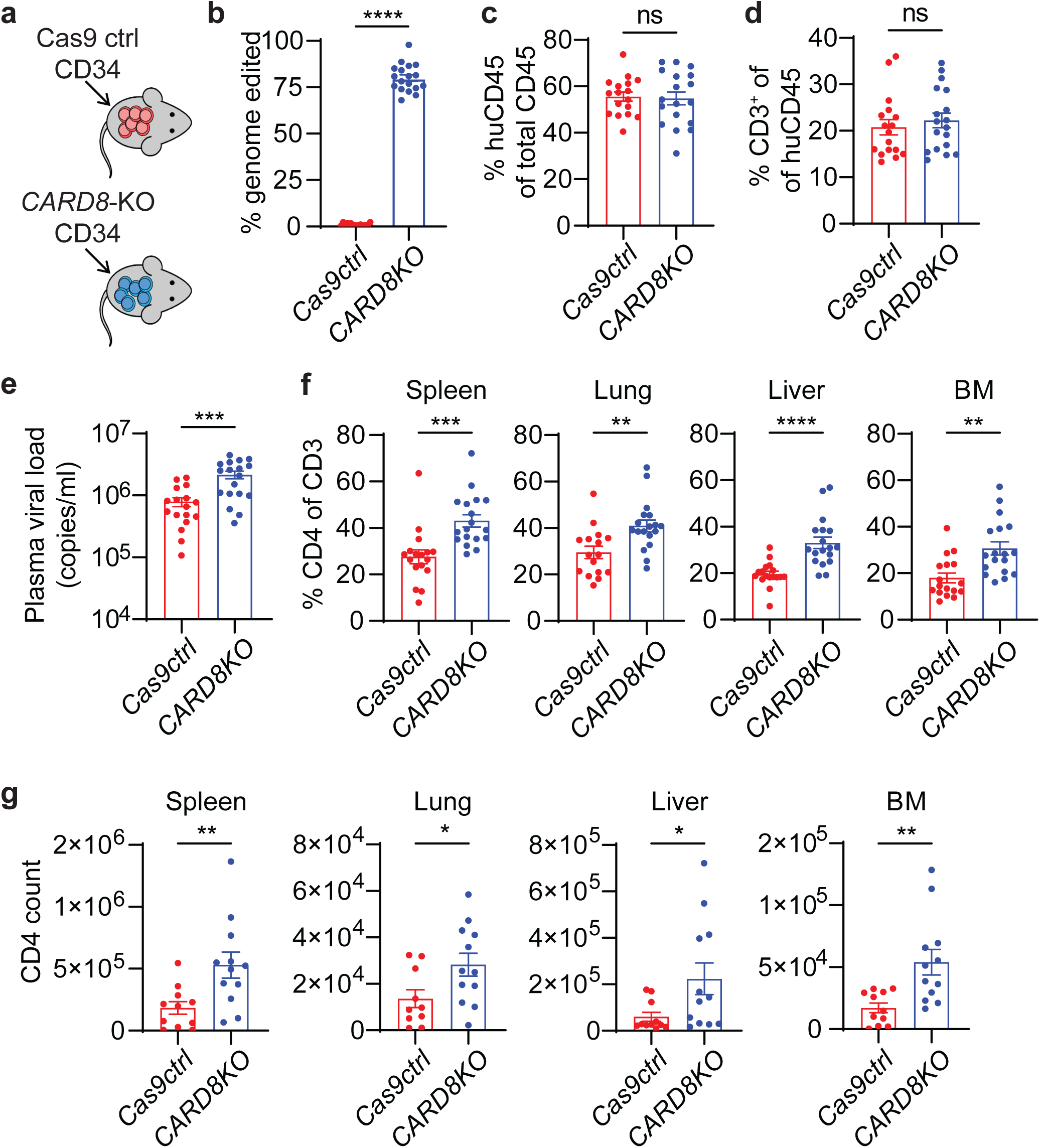
CARD8 inflammasome leads to the loss of CD4^+^ T cells in humanized mice with HIV-1 infection. **a, b**, Strategy for generating CARD8-edited humanized mice. Blood samples were collected 10 weeks after the transplantation to determine the frequency of CARD8 editing by sequencing analysis. **c, d**, Human immune cell reconstitution and T cell development in mice engrafted with *CARD8*-KO or *Cas9* control CD34^+^ cells. Blood samples were collected 10 weeks after transplantation. **e**, Plasma viral loads was measured two weeks after infection. **f, g**, The frequency and number of human CD4^+^ T cells in tissues. In b-f, data were pooled from three independent mouse cohorts. *Cas9* ctrl, N=17, except for lung (N=16); *CARD8*-KO, N=18. In **g**, data were pooled from two of the three independent mouse cohorts mentioned above. *Cas9* ctrl, N=11, except for lung (N=10); *CARD8*-KO, N=12. *p* values were calculated using an unpaired t-test. * *p* < 0.05, ** *p* < 0.01, \*\*\**p* < 0.001, and **** *p* < 0.0001. Error bars show mean values with SEM.

## Discussion

According to the UNAIDS report in 2019, only 59% of patients on ART are virally suppressed. New strategies to prevent CD4^+^ T-cell loss independent of viral suppression are urgently needed. In humans and NHPs, viral cytopathic effects contribute minimally to CD4^+^ T-cell depletion, because productive HIV/SIV infections are largely confined to activated CD4^+^ T cells, the majority of which are already destined to die rapidly regardless of the infection ^24^. Although productive infection is not required, the rapid depletion of CD4^+^ T cells by CCR5- or CXCR4-tropic chimeric simian–human immunodeficiency virus (SHIV) are confined to CCR5^+^ memory cells or CXCR4^+^ cells, respectively, strongly suggesting that the mechanism for CD4^+^ T-cell depletion involves viral entry ^9^. Since the vast majority of CD4^+^ T cells die of pyroptosis in pathogenic SIV infection in rhesus macaques ^12,13^, it is important to understand the mechanism by which HIV/SIV triggers entry-dependent inflammasome activation in CD4^+^ T cells. Using both cell-to-cell and cell-free models, we demonstrated that the viral protease encapsulated in the incoming viral particles activated the CARD8 inflammasome immediately after viral entry. We then showed that genetic ablation of CARD8 blocked the rapid CD4^+^ T cell depletion post-viral entry. Our study also highlights how resting CD4^+^ T cells are selectively targeted by HIV-1 for rapid CARD8 activation.

Furthermore, we reconstituted *CARD8*-KO or *Cas9* control human immune systems in immunodeficient mice and demonstrated that CARD8 accelerated the loss of CD4^+^ T cells in vivo. We acknowledge that CARD8 ablation only partially prevented CD4^+^ T cell destruction, which might be due to incomplete genome editing, limitations of humanized mouse models, or possible involvement of other cell death mechanisms. In addition, T cell renewal after depletion in humanized mice is less efficient, resulting in the lack of partial CD4 recovery. However, the lack of complete protection of CD4^+^ T cells in the *CARD8*-deficient immune system could also be attributed to the activation of other inflammasome sensors by HIV-1. For example, IFI16 and NLRP3 have been implicated in HIV-1-induced CD4^+^ T cell pyroptosis ^25,26^, but these two inflammasomes are only functional in certain subsets of CD4^+^ T cells and their ability to drive pyroptosis of human T cells has not been validated ^27,28^. Although the function of various components of inflammasomes has been studied in T cells, the CARD8 inflammasome is best characterized to trigger pyroptosis in human lymphocytes ^29^. In conclusion, our study reveals the role of the CARD8 inflammasome in CD4^+^ T-cell depletion by HIV-1, although its contribution to chronic inflammation is yet to be determined. Our discovery lays the foundation for CARD8-specific therapies to prevent AIDS progression.

## Acknowledgments

The following reagents were obtained through the AIDS Research and Reference Reagent Program, Division of AIDS, NIAID, NIH: ACH-2 cell line, lopinavir, T20, tenofovir, raltegravir, pNL4-3, pNL4-3-GFP.

## Author contributions

L.S. and Q.W. designed the study, analyzed the data, and wrote the manuscript. Q.W. performed all experiments.

## Data and materials availability

All data are available in the main text or the supplementary materials.

## Materials and Methods

### Mice Strain

NSG-SGM3 mice were initially purchased from Jackson Laboratory (Strain #013062) and the colony was maintained at Washington University School of Medicine. Both male and female mice were included. All animal experiments were approved by the Institutional Animal Care and Use Committee of Washington University School of Medicine (approval #20-0224).

### Human samples

Anonymous human cord blood samples were collected at the St. Louis Cord Blood Bank or the Cleveland Cord Blood Center. Anonymous peripheral blood samples were acquired from the Mississippi Valley Regional Blood Center as waste cellular products. Human tonsils were collected from elective tonsillectomies from Children’s Hospital in Saint Louis, which were provided as surgical waste without identifying information.

### Antibodies and chemicals

For Western Blot, primary antibodies against the following proteins were used: CARD8 (Abcam # ab24186), CASP1 (Abcam ab179515), NLRP1 (BioLegend #679802), ASC (Cell Signaling Technology #13833), GSDMD (Novus Biologicals #NBP2-33422) and Beta-actin (Invitrogen #MA1-140). Secondary antibodies, including horseradish peroxidase (HRP)-conjugated goat anti-mouse IgG (#7076S) and HRP-conjugated goat anti-rabbit IgG (#7074S), were purchased from Cell Signaling Technology. For flow cytometry analysis, the following fluorescence-conjugated antibodies were used in this study: Human TruStain FcX (BioLegend #422302), TruStain FcX™ (anti-mouse CD16/32, BioLegend #101320), anti-mouse CD45-FITC (BioLegend #103108), anti-human CD45-Pacific Blue (BioLegend # 304029), anti-human CD3-BV650 (BioLegend # 317324), anti-human CD4-PECy7 (BioLegend #317414), anti-human CD8-BUV395 (BD Biosciences #563795), anti-human CD45RO-PE (BioLegend #304206), anti-human CCR7-Alexa 647 (BioLegend #353218), anti-p24-PE (Beckman Coulter #KC57-RD1), and FLICA660-Caspase 1 reagent (ImmunoChemistry Technologies #9122). Other antibodies used in this study: anti-CD3 (BioLegend #300333) and anti-CD28 (BioLegend #302943). Cytokines used in this study: IL-2 (BioLegend #589106), IL-7 (BioLegend #581904), IL-15 (BioLegend #570306), SCF (STEMCELL Technologies #78062), FLT3 (STEMCELL Technologies #78009), and TPO (STEMCELL Technologies #78210). ARVs including T20, tenofovir (TFV), raltegravir (RAL), and lopinavir (LPV) were obtained from the NIH AIDS Research Program. Other chemicals used in this study: Plerixafor (AMD3100, Selleck Chem # S8030), VX765 (InvivoGen #tlrl-vad), MG132 (Cayman Chemical #10012628), Val-boroPro (VbP, Med Chem Express # HY-13233A), 1G244 (Med Chem Express #HY-116304), and PMA (Sigma #P1585).

### Plasmids and viruses

To prepare replication-defective HIV-1 reporter viruses, HEK 293T cells were transfected with pNL4-3-ΔEnv-EGFP (AIDS reagent program #11100), pNL4-3-ΔEnv-EGFP-RT-D110A-D185A or pNL4-3-ΔEnv-EGFP-IN-D116A, and the NL4-3 envelope expressing plasmid or pVSV-G. The replication-competent HIV-1_NL4-3_ was prepared by transfecting HEK293T with pNL4-3. The replication-competent HIV-1_BaL_ was produced by infecting CD8-depleted PHA-stimulated PBMCs. The culture supernatant was collected six to nine days post-infection. The lentiviruses were also produced in HEK 293T cells by co-transfecting pLKO.1puro (Addgene #8453), psPAX2 (Addgene #12260), and NL4-3 envelope expressing plasmid. Viral stocks were concentrated with the Lenti-X Concentrator (TaKaRa #631232).

### Cell culture

HEK 293T cells (ATCC #CRL-3216) and ACH-2 cells (AIDS Reagent Program #349) were cultured in DMEM or RPMI containing 10% heat-inactivated fetal bovine serum (FBS), 1 U/ml penicillin, and 100□mg/ml streptomycin (Gibco). THP-1 cells carrying doxycycline-inducible CARD8 expression cassettes were described previously ^23^ and cultured in RPMI 1640 medium supplemented with 10% FBS, 1 U/ml penicillin, and 100Cmg/ml streptomycin. Blood or tonsil CD4^+^ T cells were isolated using a human CD4^+^ T cell isolation kit (BioLegend #480010). Purified CD4^+^ T cells were used without stimulation or co-stimulated with plate-bound CD3 and soluble CD28 antibodies in the presence of 20 ng/ml IL-2 for three days. Human CD34^+^ cells were isolated from cord blood using EasySep™ human cord blood CD34 positive selection kit II (STEMCELL Technologies #17896), which were then cryopreserved in Iscove’s Modified Dulbecco’s Medium (IMDM, Sigma #S0192) containing 7.5% DMSO. The cryopreserved CD34^+^ cells were thawed and cultured in IMDM supplemented with 50 ng/ml SCF (Stemcell, #78062), 50 ng/ml Flt3L (Stemcell, #78009), and 50 ng/ml TPO (Stemcell, #78210) for 12 hours before electroporation.

### Immunoblotting and ELISA

For GSDMD cleavage detection, the unstimulated CD4^+^ T cells were spinoculated with X4-NL4-3-ΔEnv-EGFP reporter virus for two hours and incubated for another hour. For doxycycline-inducible expression CARD8^WT^, CARD8^S297A^, and CARD8^F59GF60G^ in *CARD8*-KO THP-1 cells, cells were pre-treated with doxycycline (1□μg/ml) for two days and then treated with DMSO or indicated inhibitors for 30 minutes. The treated cells were then infected with VSVG pseudotyped NL4-3-ΔEnv-EGFP viruses for three hours. Doxycycline-treated THP-1 cells were stimulated with lipopolysaccharide (50 ng/ml) for three hours, washed twice with PBS, and infected with VSVG pseudotyped NL4-3-ΔEnv-EGFP viruses for six hours. The culture supernatant was used for IL-1β ELISA (BioLegend #437004).

### Cell viability and LDH release assay

For the assessment of the cell viability, cells were washed with PBS and incubated with Zombie Violet (BioLegend #423114) for 30 minutes in accordance with the manufacturer’s instructions. The supernatants were collected after infection to determine the activity of LDH, and the LDH assay was performed according to the manufacturer’s instructions (Invitrogen #C20301).

### HIV-1 fusion assay

A reporter plasmid BLaM-Vpr was used to quantify the entry of the viruses into the cells ^30^. After pre-treating CD4^+^ T cells with different drugs for 30 minutes, the cells were infected with virions containing BLaM-Vpr for fourChours. The cells were washed in a CO2-independent medium (Thermo Fisher Scientific #18045088) and then incubated for one hour at room temperature with 100 μl CCF2/AM dye (Thermo Fisher Scientific #K1032). Afterwards, the cells were washed in RPMI and incubated overnight in a CO2-independent medium containing 10% FBS at room temperature. Flow cytometry was used to detect the green-to-blue color change as an indication of viral entry.

### Co-culture for CD4^+^ T-cell depletion

Replication competent HIV_NL4-3_ and HIV_BaL_ were used for co-culture experiments. Activated CD4^+^ T cells were infected with HIV_NL4-3_ or HIV_BaL_ for three days to reach 20%-30% infection by intracellular p24 staining. HIV-1-infected CD4^+^ T cells were used as effector cells. A total of 2 × 10^5^ donor-matched PBMCs or ToMCs were labeled with CFSE (BioLegend #423801), pre-treated with different inhibitors for 30 minutes, and then co-cultured with 2 × 10^5^ HIV-1-infected cells for six hours in a U-bottom 96-well plate without any cytokines. Flow cytometry was used to analyze the CD4 to CD8 ratio and the percentage of CCR5-positive cells within CFSE-positive cells. For ACH-2 and PBMC co-culture, 5 × 10^5^ ACH-2 cells were stimulated with 50 ng/ml PMA for 24 hours as effector cells. A total of 2 × 10^5^ PBMCs were labeled with CFSE (target cells) and treated with different inhibitors and then co-cultured with PMA-stimulated ACH-2 cells for six hours in a U-bottom 96-well plate. The CD4 and CD8 ratios of the harvested cells were determined by flow cytometry.

### Exposure to cell-free HIV-1 virions

For the exposure to cell-free HIV-1 virions, replication-defective HIV-1 reporter virus NL4-3-ΔEnv-GFP pseudotyped with the NL4-3 envelope or VSV-G was used for CD4^+^ T cells or THP-1 cells, respectively. Unstimulated CD4^+^ T cells or THP-1 cells were pre-treated with different ARVs or other inhibitors for 30 minutes and then were exposed to viruses in the 96-U-bottom plate by two hours of spinoculation and four hours of incubation before flow cytometry analysis and LDH measurement. 1000 ng HIVp24 per million cells was used unless noted otherwise. The following inhibitor concentrations were used: AMD3100 5 μM, T20 5 μM, TFV 1 μM, RAL 5 μM, LPV 5 μM, VX765 50 μM, MG132 10 μM, and VbP 5 μM. 1G244 was used at 500 nM unless noted otherwise.

### CRISPR knockout in CD4^+^ T cells and CD34^+^ cells

Cas9-sgRNA ribonucleoprotein (RNP) complexes were electroporated into CD4^+^ T cells and CD34^+^ cells using the Lonza 4D Nucleofector system. The recombinant Cas9 protein was obtained from IDT (#1081059). The modified synthetic sgRNAs were purchased from Synthego and the sequences are listed in **Supplementary Table 1**. RNP complexes were prepared by mixing Cas9 (40 pmol) with sgRNA (100 pmol) and incubating them for 10 minutes at room temperature. 2 ×□10^6^ unstimulated CD4^+^ T cells or 0.2 × 10^6^ CD34^+^ cells were washed with PBS and resuspended in 20□μl buffer P3 (Lonza #V4XP-3032). To deliver two specific sgRNAs targeting one gene, 2□μl of each RNP complex were then mixed with the cell suspension and transferred into a 16-well reaction cuvette of the 4D-Nucleofector System (Lonza #V4XP-3032). The CD4^+^ T cells and CD34^+^ cells were electroplated using the programs EH-100 and DZ-100, respectively. After electroporation, CD4^+^ T cells and CD34^+^ cells were resuspended in 100 μl of prewarmed RPMI 1640 or IMDM and transferred to a 96-well plate to recover for 30 minutes at 37°C. The electroporated CD4^+^ T cells were cultured for three to four weeks under low concentrations of human IL-7 (2 ng/mL) and IL-15 (2 ng/mL) before *in vitro* testing. The electroporated CD34^+^ cells were then washed with PBS before being transplanted into the mice.

### Flow cytometry analysis

The mouse tissue cell suspensions were first incubated with Zombie NIR (BioLegend #423106) in PBS for 30 minutes at 4°C, followed by an incubation with Human TruStain FcX and TruStain FcX (anti-mouse CD16/32) blocking antibodies for 10 min at room temperature. The cell suspensions were then incubated at 4°C with fluorescence-conjugated antibodies for 30 minutes to stain surface antigens. In all experiments, stained cells were acquired on a BD LSR Fortessa, X20, or Accuri C6 (BD Biosciences), and data were analyzed by the FlowJo software.

### Generation and HIV-1 infection of humanized mice

One-to three-day old newborn NSG-SGM3 mice were preconditioned with sublethal irradiation (100 cGy) followed by an intrahepatic injection of 3-4×10^4^ unmodified or Cas9-edited CD34^+^ cells. Reconstitution of human CD45^+^, CD3^+^, and CD4^+^ T cells was assessed 10 to 12 weeks after engraftment. Experimental groups were assigned randomly. Engrafted mice were infected with HIV_BaL_ (10 ng p24/mouse) through retro-orbital injection. Blood samples were collected by retro-orbital or submandibular vein bleeding to quantify plasma HIV-1 RNA. The plasma viral RNA was extracted using a Quick-RNA Viral Kit (Zymo Research #R1035) and reverse transcribed using the ProtoScript II Reverse Transcriptase (NEB #M0368). A ten-fold serial dilution of HIV-1 genomic DNA served as a standard for measuring plasma viral RNA by the HIV-1 gag-based qPCR assay ^31^.

### Amplicon deep sequencing

In the first step, corresponding primers were used for the CARD8 loci (CARD8-F: GCTGCTTAAATCCAAGTCCTGGGG; CARD8-R: CTGGTGGGCGGCCCCTTG). In a second round of PCR using primers (**Supplementary Table 2**) containing sample-specific barcodes and adapters, amplicons were sequenced for 2□×□150 paired-end reads with MiSeq Sequencing (Illumina). The CRISPResso software was used to analyze the deep sequencing data ^32^.

### Statistical Analysis

The statistical analysis was conducted using Prism 9 (GraphPad). The methods for statistical analysis are described in the figure legends. The error bars indicate the standard error of the mean.

